# Queen loss fails to elicit physiological and transcriptional responses in workers of the invasive garden ant *Lasius neglectus*

**DOI:** 10.1101/2024.01.26.577224

**Authors:** Megha Majoe, Nicole Stolarek, Joel Vizueta, Zijun Xiong, Lukas Schrader, Jacobus J. Boomsma, Susanne Foitzik, Romain Libbrecht, Volker Nehring

**Affiliations:** Institute of Organismic and Molecular Evolution, Johannes Gutenberg University Mainz, Germany; Department of Evolutionary Biology and Ecology Institute of Biology I (Zoology), Albert Ludwig University of Freiburg, Germany; Villum Centre for Biodiversity Genomics, Section for Ecology and Evolution, Department of Biology, University of Copenhagen, Denmark; BGI Research, Wuhan 430074, China; College of Life Sciences, University of Chinese Academy of Sciences, Beijing, China; Institute for Evolution and Biodiversity, University of Münster, Germany; Section for Ecology and Evolution, Department of Biology, University of Copenhagen, Denmark; Insect Biology Research Institute, UMR 7261, CNRS, University of Tours, France

**Keywords:** worker sterility, gene expression, fat body, major transitions in evolution, germline, soma, reproductive division of labour

## Abstract

Insect colonies with morphologically distinct castes have been called superorganisms because their queens and workers are functionally analogous to the germline and soma in metazoan bodies. In the haplodiploid ants, workers typically lost the sperm storage organ but retained ovaries so they can lay unfertilized eggs. Worker reproduction often occurs after queen loss and is accompanied by a number of physiological changes. However, in some evolutionarily derived ants, workers have become functionally sterile and in many of these species colonies contain multiple queens and can readily raise replacement queens - a syndrome characteristic for invasive ants. We hypothesized that the combination of full worker sterility and regular queen replacement should have eliminated important aspects of the physiological interdependence between queens and workers. We tested this hypothesis by analysing fat body gene expression and worker resistance to oxidative stress in colonies of the invasive ant *Lasius neglectus* with and without queens. We found age-related transcriptional shifts between young and old queens and young and old workers, suggesting rapid ageing in all castes. However, the removal of any queens in controlled experiments failed to elicit changes in the transcriptional activity and oxidative stress resistance of workers, consistent with our hypothesis. The invasive syndrome of this ant may thus have led to a somatic work force that evolved to be physiologically independent of queen presence.

## Introduction

The social insects realized major transitions in organizational complexity when they evolved obligate division of reproductive labour between morphologically differentiated queen and worker castes. This irreversible transition to superorganismality evolved convergently in the hymenopteran corbiculate bees, vespine wasps, and ants, and in at least some of the isopteran termites (Boomsma & Gawne, 2018, Boomsma 2022, Revely et al., 2021). In the ancestors of all of these lineages, the queens became morphologically and functionally specialized as the germline of the colony, laying all or almost all of the eggs. The workers primarily adopted a variety of somatic functions including brood care, foraging, and defence. This germline-soma analogy never applies to the open societies of cooperatively breeding vertebrates, insects, and other arthropods, because most if not all helpers at the nest retain the potential to become reproductive later in life (Boomsma, 2022).

Obligate reproductive division of labour is often reflected not just in morphology but also in lifespan. In perennial superorganisms with a single inseminated queen (ants, bees, wasps) or a single royal pair (termites), the reproductives can live for decades, while the non-reproductive workers often survive for just a few months (Keller & Genoud, 1997b; Korb, 2016; Lucas & Keller, 2020; Parker, 2010). The mechanisms responsible for caste-differentiated ageing remain to be fully resolved, but it is clear that queens have upregulated gene regulatory networks for body maintenance that are instrumental for achieving longer life spans (Lin et al., 2021; Monroy Kuhn et al., 2021; Negroni et al., 2019). The life-span differences between queens and workers are less pronounced in the smaller annual superorganisms of bumblebees and yellowjacket wasps, and they have become secondarily reduced in the perennial superorganisms that have either replacement procedures for single queens or ‘polygyny’ in the form of multiple coexisting queens in the same colony (Keller & Genoud, 1997; Boomsma et al., 2014).

Despite their role as specialized colony soma, the workers of many superorganismal lineages can still reproduce. However, ovarian development in workers is often conditional on the queen’s capacity for egg-laying (Monnin, 2006), so that queens produce pheromones to signal their presence and fecundity. Structurally very similar queen pheromones have been identified in multiple social hymenopteran insects, consistent with these compounds having originated in a common solitary wasp ancestor (Grüter & Keller, 2016; Holman et al., 2010; Keller & Nonacs, 1993; Shimoji et al., 2017). Hymenopteran workers typically do not develop their ovaries as long as queen pheromones are produced, but they readily switch from a somatic to a reproductive role after the queen dies. This results in the production of male-destined haploid eggs by the workers (Khila & Abouheif, 2010; Meunier et al., 2010). Yet workers cannot replace a queen themselves because they lack the capacity to lay the fertilized eggs that could yield female workers and queens. Queen loss typically triggers a whole cascade of physiological changes in workers. It affects the expression of hundreds of genes, not only for reproduction but also for longevity-related processes such as oxidative stress management, resulting in reproductive workers often outliving their non-reproductive sisters (Amsalem et al., 2017; Giehr et al., 2020b; Kennedy et al., 2021a; Negroni et al., 2021; Seehuus et al., 2006a; Tsuji et al., 1996).

Young workers labor inside the nest and are more likely to activate their ovaries and start reproducing than older workers who tend to carry out tasks outside the nest (Dijkstra et al., 2005; Leonhardt et al., 2016; Lin et al., 1999; Nakaoka et al., 2008; Richardson et al., 2021).Retaining the potential for worker reproduction in response to queen loss is beneficial when it allows the colony to pass on more gene copies to future generations before it collapses. However, premature worker reproduction induces unproductive intra-colony conflict and may happen because the fitness interests of haplodiploid workers and queens, and among the workers individually, are not aligned. In many species, these conflicts are suppressed by policing when workers attack their own egg-laying sisters or when they destroy worker-laid eggs (Brunner & Heinze, 2009; Dijkstra et al., 2010; Helanterä & Sundström, 2007; Schmid et al., 2013; Stroeymeyt et al., 2007; Van Oystaeyen et al., 2014). However, none of this applies in the various ant lineages where worker ovaries have been lost over evolutionary time or have become otherwise non-functional. To our knowledge, the social consequences of worker ovary loss have not been systematically evaluated, but it seems clear that they should be different between lineages that retained the ancestral single-queen (i.e. monogynous) condition and lineages that secondarily evolved obligate polygyny, i.e. the coexistence and recurrent recruitment of multiple queens in each colony.

Because fully sterile workers cannot reproduce, one might expect that they should not react to the loss of a single mother queen. However, this would overlook that orphaned sterile workers could still increase their lifetime indirect fitness by raising any remaining sibling eggs and larvae into dispersing reproductives, an agenda that might well require significant changes in worker physiology. The sterile workers of *Atta* leaf-cutting ants, which have a single irreplaceable queen, are known to become more resistant to oxidative stress when they lose the queen (Majoe et al., 2021). This suggests that orphaned workers benefit from prolonging their own lives when raising the remaining sexual offspring of their late queen has become an unconditional priority. This shift in prioritizing brood care has also been documented for another leaf-cutting ant genus, *Acromyrmex* (Dijkstra & Boomsma, 2007). Yet another study indicated that when the sole colony queen of the monogyne form of the red imported fire ant *Solenopsis invicta* dies, it is the gynes (virgin queens) that start producing male offspring from unfertilized eggs (Wurm et al., 2010), while the sterile workers continue their previous tasks, consistent with queen loss having only subtle effects on whole-body transcriptomes of workers (Manfredini et al., 2014). These findings indicate that the inevitable end of a monogynous colony-life-cycle initiates a terminal bout of reproductive investment that involves alterations in the physiology and behaviour of the orphaned queen offspring.

Fully sterile workers are also a feature of the unicolonial and often invasive ants that are invariably characterized by high degrees of polygyny, low relatedness among the workers, and highly polydomous colony structures (Helanterä et al., 2009). In contrast to monogynous and weakly polygynous ant species, the loss of queens is virtually impossible in such colonies, because virgin queens are produced in large numbers, fertilised locally without the need for a dispersal flight, and then migrate freely between nests (Helanterä et al., 2009; Silverman & Brightwell, 2008; Tsutsui et al., 2000). This implies that colony survival has become decoupled from individual queen mortality, and that no phase of terminal investment is expected. We therefore hypothesized that any form of selection for maintaining physiological responses to queen loss in the worker caste must have disappeared in unicolonial ants. In addition, there should be little selection for longevity in queens or workers, as both can be easily replaced.

We tested these expectations using the invasive species *Lasius neglectus*, a unicolonial formicine ant that was discovered rather recently (Van Loon et al., 1990) and that is native to Central Asia (Stukalyuk et al., 2020). The species has been introduced to Europe via the Black Sea area (Cremer et al., 2008) and spread further across most of Europe through human-assisted dispersal (Boomsma et al., 1990; Seifert, 2000). *Lasius neglectus* is a suitable model species because workers have never been observed to lay eggs, consistent with their ovaries regressing completely within the first four months after hatching (Espadaler et al., 2004; Espadaler & Rey, 2001; Gotoh et al., 2016; Stukalyuk & Radchenko, 2018). Other *Lasius* species have workers with active ovaries (Bourke, 1988) and reproductively active *Lasius neglectus* queens still produce the same fertility pheromone as other monogynous *Lasius* ants (Holman et al., 2013). This suggests that a possible decoupling of worker physiology from individual queen mortality is evolutionary recent, consistent with the existence of a sibling species, *Lasius turcicus*, which has not become invasive and unicolonial (Cremer et al., 2008).

We investigated whether queen removal affected unicolonial *Lasius neglectus* workers in a similar way as one would expect it to affect workers of single-queen species (Alaux et al 2006; Amsalem et al., 2017; Giehr et al., 2020b; Kennedy et al., 2021a; Negroni et al., 2021; Seehuus et al., 2006a; Tsuji et al., 1996; Manfredini et al., 2014, Majoe et al 2021). We examined the molecular physiology of workers focusing on the transcriptomes of the fat body, an important metabolic organ analogous to the human liver and adipose tissue, which is also involved in regulating fertility (Partridge et al., 2011; Tatar et al., 2003; Yan et al., 2022). We also tested for effects on oxidative stress resistance, a response to queen loss previously found in workers of other ant species (Majoe et al., 2021) and the honey bee (Kennedy et al., 2021; Seehuus et al., 2006). We examined workers collected inside laboratory nests close to the brood pile, and outside the nest in the foraging arena, as respective representatives for young nurses and older foragers. Finally, we obtained transcriptomes of young and old queens. Consistent with our main hypothesis, we found that queen presence/absence had no effect on workers’ susceptibility to oxidative stress or on their molecular physiology. However, we observed strong differences between ‘inside-workers’ and ‘outside-workers’, both in terms of overall gene expression profiles and oxidative stress susceptibility, consistent with rapid ageing in these obligately sterile workers.

## Material and Methods

### Study site, collection, and laboratory maintenance

All ant colonies used for this study were collected from the Jena Botanical Garden in Germany on three collecting trips in July 2019, September 2019, and July 2020. Alates (winged virgin reproductives), brood, workers, and queens were found close to the surface near flowerbeds and in sand piles in disturbed, slightly damp areas of the garden. We assigned ants collected from different sand piles or areas that were about 5-10 m apart to independent ‘replicate colonies’ and kept them separate. The ants could place their brood and queens in glass tubes half-filled with water and plugged with cotton wool. These tubes were covered with aluminium foil to provide a darkened environment and placed in Fluon™-coated boxes (20 x 20 x 5 cm), which acted as foraging arenas. The ants could access an agarose-based jelly-mixture containing honey and egg ad libitum for food. All boxes were placed in a climate cabinet with a temperature set to 25°C and a 12h light:12 h dark cycle.

### Creating subcolonies

Dealate queens collected within field colonies were later assigned to be ‘old’ queens in our transcriptome study. ‘Young’ queens (in fact, gynes at the time) were those collected with wings and thus presumably still uninseminated. We separated the dealate old queens and winged gynes arising from the same replicate colonies and set each group up with half of the workers from that colony. Then we added males to the sub-colonies with the winged gynes, so they could become mate. Once inseminated, these young queens typically shed their wings and became reproductive.

To investigate the effect of queen removal on workers, we established queenless subcolonies from the young-queen and old-queen colonies, which were composed of about 80 workers and larvae. These subcolonies were kept for four weeks, which is sufficient in other ant species to elicit ovary development in workers (Holman et al 2013; Kohlmeier et al., 2017). Before sampling the workers, we starved them for three days to increase foraging activity. We then marked individuals that were the first to arrive at new food on two consecutive days as outside/foraging workers and those that stayed close to the brood pile as inside/nursing workers.

### Oxidative stress experiment with workers

The marked workers and some brood from each subcolony (with or without a queen) were used to produce yet another nested level of small paired subcolonies consisting of five outside workers and five inside workers. Each of these was randomly assigned to be a control or treatment subcolony. Over the next twelve days, 196 workers from five such colonies were subjected to paraquat, which induced oxidative stress, or water as a control treatment. Every two days, the paraquat or water solution was applied with a brush on the head of each ant, followed by a period of 2 hours of isolation allowing each ant to self-groom and ingest the solution. Worker survival was then tracked every day. The treatment and observation of survival followed the protocol of a previous study on susceptibility of myrmicine ant workers to oxidative stress (Majoe et al., 2021).

### Transcriptome analyses

Six weeks after the creation of the queenless subcolonies, we sacrificed workers from queenless and queenright subcolonies and dissected their fat bodies within a time window of four hours (Supplementary Table 1). We pooled the fat bodies of four workers into a single sample and stored samples in Trizol at −80°C. The queens were sacrificed in the same month (October 2020) but fat bodies were large enough to keep them as individual queen samples (Supplementary Table 2). At the time of sampling, the young queens of the five colonies were about six months old. Since *L. neglectus* colonies in the field have been observed to produce winged sexuals only once a year (Espadaler & Rey, 2001; Stukalyuk et al., 2020; Stukalyuk & Radchenko, 2018), we inferred that the old queens were at least one year or older than young queens when we sacrificed them. We also sacrificed two queens that were 2.5 years or older and obtained their fat bodies. We confirmed that all queens had filled spermathecae and measured the ovariole lengths of a subset of five old and five young queens. There was no indication that ovariole lengths were related to age (n = 10; χ^2^=0.02, p=0.89; Supplementary Fig 1).

### Genome assembly and annotation

*Lasius neglectus* specimens, which were collected in 2015 in Jena and since then reared in the laboratory, were provided by the Sylvia Cremer group and used for genome sequencing as part of the Global Ant Genomics Alliance (GAGA) initiative (Boomsma et al., 2017). High-quality DNA was extracted from workers using a phenol/chloroform phase separation extraction protocol. Single-tube long fragment read (stLFR) libraries were prepared and sequenced by BGI using the BGISEQ-500 platform. In addition, total RNA from whole individual queens, males, and brood was extracted using the RNeasy Plus Mini Kit from QIAGEN. The quality of extracted RNA was assessed with the Agilent 2100 Bioanalyzer. RNA-seq libraries were prepared with >1µg total RNA using the MGIEasy RNA Library Prep Kit (BGI) and sequenced on the BGISEQ-500 platform.

Raw stLFR reads were first cleaned from adaptors and PCR duplicates, after which the barcode IDs were assigned in the read names using the stlfr2supernova_pipeline (https://github.com/BGI-Qingdao/stlfr2supernova_pipeline). The clean reads were then assembled using MaSuRCA v3.3.0 (Zimin et al., 2013); “JF_SIZE = 2500000000”) and the resulting assembly was further scaffolded using the barcoding information from stLFR reads to assemble contigs into scaffolds with the SLR-superscaffolder pipeline (https://github.com/BGI-Qingdao/SLR-superscaffolder). Putative duplicated scaffolds were identified and filtered using Funannotate “clean” pipeline v1.8.3 (Palmer & Stajich, 2022). The genome assembly was screened for putative contaminations from other organisms using a pipeline established and optimized for the ant genomes in the GAGA project. In brief, we compiled separate databases containing 1908 complete bacterial genome sequences, 43 complete insect genome sequences, as well as databases containing corresponding bacterial or insect CDS sequences. We divided the *L. neglectus* genome assembly in 2000 bp sliding windows (with 500 bp overlap) and searched each window against the different insect and bacterial databases using mmseqs (release_12-113e3) and identified the single best hit (according to bitscore) for each sliding window against each database. For each scaffold, we calculated the ratio of windows showing higher similarity to bacterial than to eukaryotic databases and used this, along with coverage and GC content information, as three lines of independent evidence to identify contaminant scaffolds. Genome assembly quality was then evaluated using contiguity metrics, gene completeness with BUSCO v5.1.2 (Manni et al., 2021), and consensus quality (QV) and k-mer completeness using Merqury (Rhie et al., 2020; Supplementary Table 3). Finally, some mitochondrial (*CO1* and *CytB*) and autosomal gene markers (*Wingless*, *LwRh*, *AbdA*, *ArgK*) were annotated in the genome assembly using BITACORA (Vizueta et al., 2020) to confirm the species identity from *Lasius neglectus* sequencing data.

Homology-based and *de novo* methods were used in combination to identify the transposable elements (TEs). The genome sequences were aligned against the Rebase TE library (v25.03) (Roberts et al., 2015) and the TE protein database using RepeatMasker and RepeatProteinMask (version 4.1.2) (Smit et al., 2013). In addition, RepeatModeler v2.0.2 (Smit & Hubley, 2021) was used to build a *de novo L. neglectus* repeat library, which was subsequently used to annotate repeats using RepeatMasker. TRF v4.10.0 (Benson, 1999) was then used to find tandem repeats with parameters: "Match = 2, Mismatch = 7, Delta = 7, PM = 80, PI = 10, Minscore = 50". Finally, we combined all evidence resulting in 26.8% of the assembled genome being repeat sequences, with a total length of 68 Mb (Supplementary Table 3).

General genome annotation was conducted by combining gene annotation from several sources using a pipeline optimized for the ant genomes generated by the GAGA project. First, publicly available RNA-seq data from *L. neglectus* workers and queens (NCBI Bioproject PRJDB4088, Morandin et al., 2016), as well as the GAGA generated RNA-seq data were aligned to the reference repeat soft-masked genome assembly using the STAR v2.7.2b default options (Dobin et al., 2013). In addition, we retrieved the publicly available gene annotations from the fruit fly *Drosophila melanogaster*, the red flour beetle *Tribolium castaneum*, the parasitoid wasp *Nasonia vitripennis*, the honeybee *Apis mellifera*, the clonal raider ant *Ooceraea biroi* and the Florida carpenter ant *Camponotus floridanus* (Hoskins et al., 2015; Kim et al., 2010; McKenzie & Kronauer, 2018; Richards et al., 2008; Shields et al., 2018; Weinstock et al., 2006; Werren et al., 2010). The annotations from these insect species were used to conduct homology-based gene predictions using GeMoMa v1.7.1, which also incorporated the RNA-seq evidence for splice site prediction (Keilwagen et al., 2019). Second, the independent RNA-seq alignments were merged creating a consensus GTF (Gene transfer format) using Stringtie v2.1.5 (Kovaka et al., 2019), and BestORF (Molquest package, Softberry) was used to identify open reading frames (ORF) in the transcript sequences, after which the transcripts with incomplete ORFs were filtered out. Third, we randomly selected ∼1,000 high–quality genes from the GeMoMa prediction to train Augustus v3.2.2 (Stanke et al., 2008). *De novo* gene prediction was then performed using Augustus with the repeat-masked genome, filtering out genes with lower length than 150 bp or incomplete ORFs. Finally, gene annotations from the three independent sources of evidence were combined, generating a final gene annotation for the *L. neglectus* genome, from which we also generated an annotation with a single representative isoform per gene (i.e.: the longest isoform was kept as the representative one). Transposon-related proteins were identified and filtered using a BLASTP search against the Swissprot database and the transposable element protein database from RepeatMasker. The quality of the genome annotation was finally evaluated in terms of completeness based on the BUSCO Hymenoptera dataset (Supplementary Table 3).

### RNA extraction and analysis

RNA from queens and workers was extracted using a Trizol: Chloroform: Isoamylacohol mixture based on an in-house protocol (adapted from Lin, Werle, & Korb, 2021) followed by cleaning steps employing the Qiagen RN-easy Mini Kit. The concentration and quality of the isolated RNA was checked on an Agilent Bioanalyzer (Agilent RNA 6000 Nano Kit) at the Beijing Genomics Institute (BGI). The libraries were prepared and sequenced by BGI using the Illumina HiseqXTen sequencing platform, yielding 150 bp paired-end reads with a sequencing depth of 23.85 +/-1.97 (mean +/-sd) million reads.

Raw reads for 47 samples (9 old queens, 9 young queens, 8 queenright old worker pools, 8 queenright young worker pools, 6 queenless old worker pools, 7 queenless young worker pools; Supplementary Tables 1 & 2) were trimmed using fastp (version 0.2), such that all reads that passed had a minimum length of 70 bp. The trimmed results were checked with MultiQC-1.7 and the quality of the reads was assessed with FastQC (version 0.11.8). HISAT2 (version 2.1.0) was used to align the trimmed sequences to the *L. neglectus* genome. All samples were assigned to the genome with an alignment rate of 83.7-91.66%. The SAM files generated by HISAT2 were first converted to BAM files and then sorted by name using SAMtools (version 1.9). The sorted BAM files were used to create gene count tables with Htseq-count (Htseq version 0.11.2; settings: -f bam, -i ID, -t gene, -m union, -r name, --stranded=no) after which a single read count table consisting of all samples was compiled. In addition, a metadata table was assembled that contained information about each sample. The data were further analyzed using R-Studio (R version 4.0.4) and we used the DESeq2 package (version 1.30.1) to analyse differential gene expression. Since we aimed to explore different predictors for queens and workers, we decided to analyse the data sets for the two castes separately. In each dataset, we removed genes that had low read counts in more samples than could potentially be expected even if only one of the experimental groups would express the gene. We thus only kept genes with at least ten reads in at least five worker samples (out of 14059 genes in the genome, 10691 remained) or at least eight queen samples (10022 genes remained out of 14059).

For the queen analysis, we used "age" (young vs. old) and "replicate colony" as factors and determined the effect of age using the Likelihood Ratio Test (LRT), while controlling for the effect of replicate colony. The worker-specific model included the factors "replicate colony", "queen presence/absence", "worker position” (inside/outside), and the interaction between the latter two variables. Since the interaction did not reveal any differentially expressed genes (DEGs), we built an additive model with the three factors and used the LRT to test the effects of location and queen presence separately, while controlling for the other two factors. Adjusted p-values of 0.05 were used as a cut-off to obtain all genes whose expression was significantly explained by the factor of interest. The p-values were always corrected for false discovery rates using the Benjamini-Hochberg procedure, yielding ‘adjusted p-values’.

### Gene annotation, functional enrichment and comparison to known candidate gene lists

We ran Interproscan (v5.54) on the *Lasius neglectus* protein coding genes to obtain functional annotations and process GO (Gene Ontology) IDs associated with our differentially expressed gene (DEG) lists. We extracted proteins associated with age and position from the queen and worker analyses respectively. The GO terms from the Interproscan results were used to perform enrichment analyses using topGO (v 2.4.2) on R-Studio (R version 4.0.5). We used the ’weight01’ algorithm to determine the nodes and Fisher’s test with p < 0.05 after Benjamini Hochberg correction to test which GO terms were enriched in our DEG lists on a background of the 8255 unique proteins in the *L. neglectus* proteome that had GO IDs associated with them. We ran a local BLASTp (BLAST v 2.5.0+) of the entire proteome against a database of nine insect species: *Drosophila melanogaster, Apis mellifera, Lasius niger, Linepithema humile, Solenopsis invicta, Ooceraea biroi, Acromyrmex echinatior, Atta colombica* and *Temnothorax curvispinosus*. We chose the hit with the lowest bit score among the ten hits with the lowest e-values that had e-values of e^-5^ or lower. The functional annotation of genes associated with specific GO terms could then be determined by retrieving the selected *Lasius neglectus* genes from the full list (Supplementary Table 4).

We also specifically investigated some of the candidate genes using Orthofinder (v.2.5.4) to obtain orthogroups of similar *L. neglectus* and *D. melanogaster* proteins. We then extracted those orthogroups associated with 387 proteins encoded by 123 candidate genes implicated in ageing in *D. melanogaster* and other social insects. Referred to as ‘TI-J-LiFe’ genes (Korb et al., 2021), this list features genes in IIS (Insulin/Insulin Like Growth factor), TOR (Target of Rapamycin), and Juvenile Hormone signalling pathways in *Drosophila melanogaster.* Similarly, we extracted orthogroups associated with 85 proteins from 33 known oxidative stress genes from *D. melanogaster* (Kramer et al., 2021).

We additionally searched specifically for the expression of any *Vitellogenin* (*Vg*) genes in the *L. neglectus* genome because of the role of *Vg* and their copies in determining fecundity, regulating oxidative stress, and division of labor. The *Vg* genes were identified by conducting a reciprocal BLASTP search of the *L. neglectus* proteome against a database of annotated Vitellogenin amino acid sequences with an e-value of 1e-5. The sequences were from the 33 species used in Kohlmeier et al. (2018).

## Results

### Worker task, but not queen presence, affects worker resistance to oxidative stress

Worker survival was significantly lower under paraquat-induced oxidative stress compared to the control treatment (Fig. 1; treatment, n = 196, χ^2^ = 26.1, p < 0.001). Foragers had lower survival overall (Fig. 1cd; inside *versus* outside, n = 196, χ^2^ = 17.6, p < 0.001). However, we did not detect an effect of queen presence on worker survival (Fig. 1ab; queen presence *versus* absence, n= 196, χ^2^ = 0.53, p = 0.46), neither for inside (nurses) nor for outside (foragers) workers (interaction queen presence/absence: inside/outside worker position, n = 196, χ^2^ = 0.19, p = 0.66). Similarly, there was no interaction between paraquat treatment/control and queen presence/absence (n = 196, χ^2^ = 1.8, p = 0.18), or between paraquat treatment/control and inside/outside position (n=196, χ^2^ = 0.32, p = 0.57).

**Figure 1:**
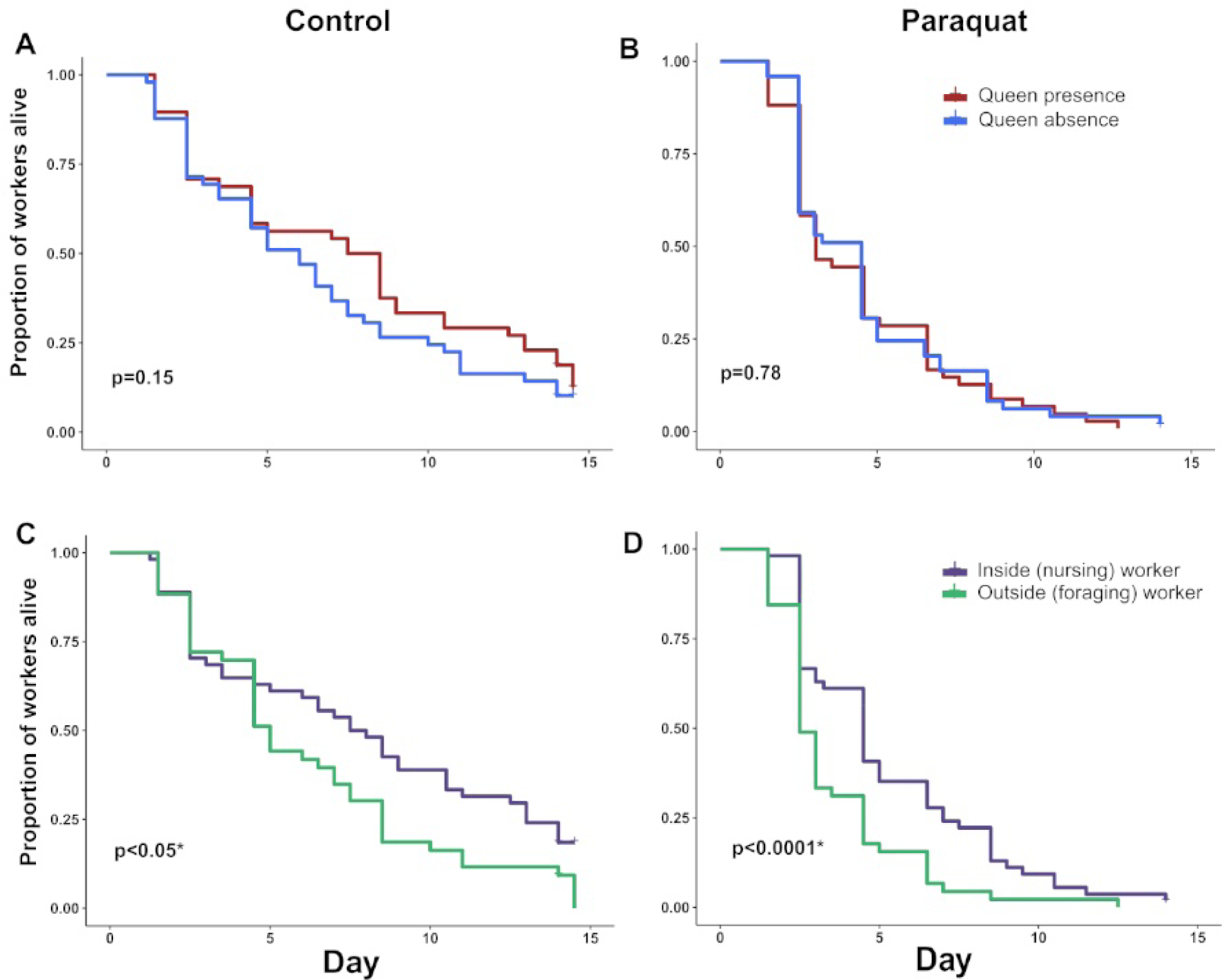
Survival of *Lasius neglectus* workers over 14 days of the experiment. Queen presence did not affect worker survival under control (a) or paraquat-induced oxidative stress conditions (b), but inside (nurse) workers tended to survive better (c,d), in particular under paraquat treatment (d). P-values shown inside the plots are from Cox regression models between the two groups depicted in the plot (n = 196 ants in total).

To make sure that we did not miss potentially weak effects of worker position or queen presence/absence because they might have been overshadowed by the strong treatment effect, we additionally ran separate models for paraquat treatment and control. Queen presence still did not affect worker survival; neither in the control (queen presence/absence, n = 97, χ^2^ = 2.1, p = 0.15), nor in the paraquat treatment (queen presence/absence, n = 99, χ^2^= 0.07, p = 0.78). There was also no interaction between queen presence and worker position on worker survival in either the control treatment (queen presence/absence: inside/outside position, n = 97, χ^2^ = 0.05, p = 0.82) or the paraquat treatment (queen presence/absence: inside/outside position, n = 99, χ^2^ = 0.63, p = 0.42). This implied that the sampling position of workers explained most of the variation in worker survival, both in the control experiment (inside/outside position, n = 97, χ^2^ = 5.95, p = 0.015) and in the paraquat-treatments (inside/outside position, n = 99, χ^2^ = 11.58 p < 0.001; Fig. 1, Supplementary Table 5).

### Gene expression analyses

Queen presence did not influence gene expression in the fat body of workers, neither for outside nor for inside workers after correcting for false discovery rate (all p_adjusted_ >0.05 for both queen presence alone and for the interaction term of queen presence and inside/outside location (Fig. 2 a,b). In contrast, we detected 2743 differentially expressed genes (DEGs) between inside and outside-workers, with 1444 of these being overexpressed in inside workers and 1229 in outside workers. This indicates that we had enough statistical power to identify differential gene expression in the inside/outside contrast, which we thus assumed was also the case when we compared the workers with and without queen.

**Figure 2:**
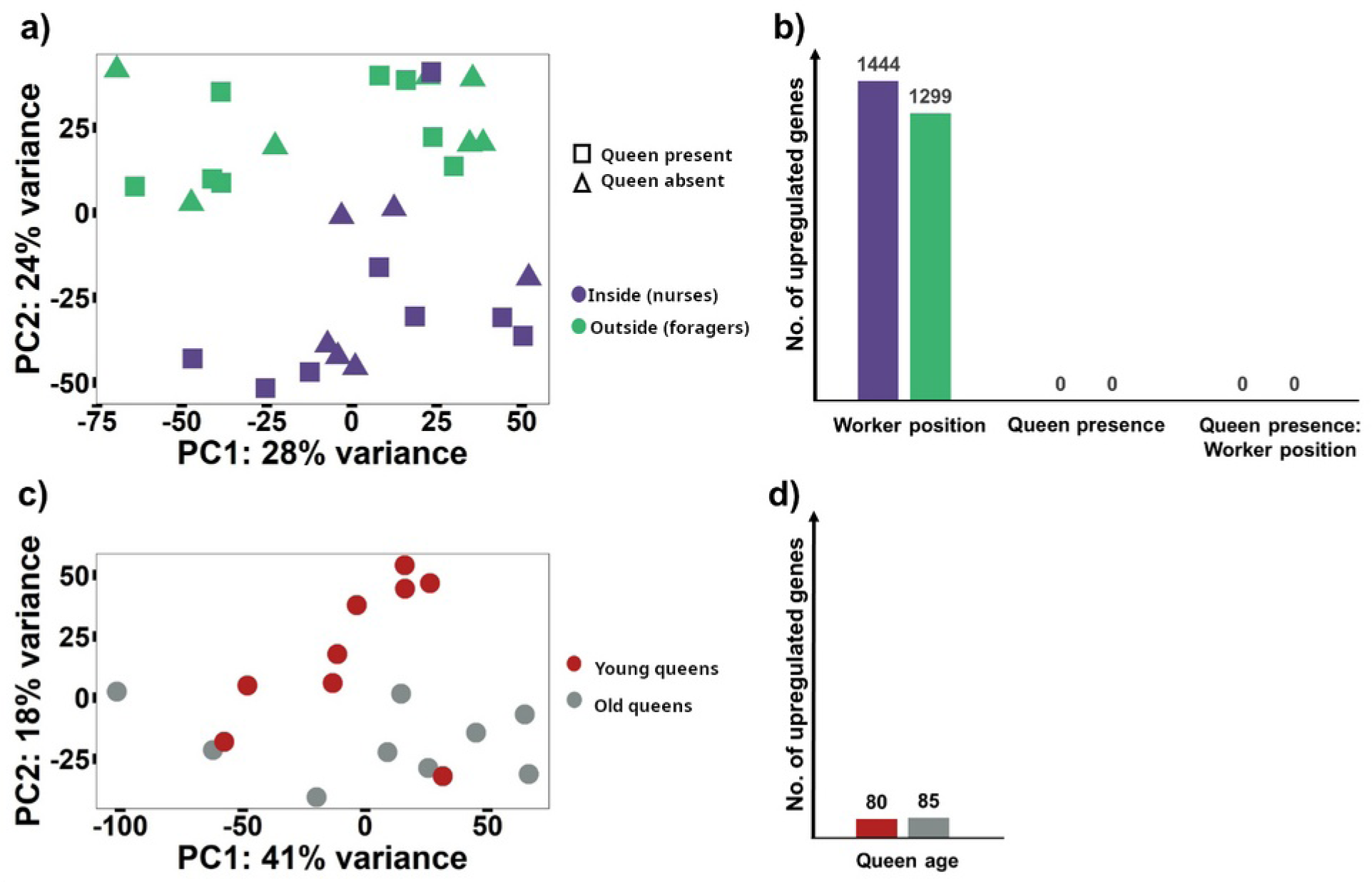
Principal component analyses based on the expression of (a) all genes expressed in workers and (c) all genes expressed in queens. (b) The expression of many genes was affected by the inside nursing versus outside foraging contrast among the sampled workers (Deseq2 padjusted < 0.05). No genes were affected in their expression by the queen presence/absence contrast alone or by its interaction with nursing/foraging. (d) Queen age also affected gene expression, but the number of differentially expressed genes was comparably low.

DESeq2 analysis of the queen samples identified 165 genes that were differentially expressed between young and old queens (Fig 2c,d, Supplementary Table 5). Of these, 80 genes were significantly overexpressed in younger queens while 85 genes were significantly overexpressed in older queens (Fig 2d). There is thus an order of magnitude difference in DEGs between young and old queens compared to young, inside, and old, outside workers.

Six GO terms were significantly enriched for workers and four for queens (p < 0.05 after correcting for multiple testing; Figure 3; Supplementary Table 6). The genes upregulated in young, inside workers were related to DNA replication (Figure 3a, Supplementary Table 5). Interestingly, the GO term ‘proteolysis involved in cellular protein catabolic processes’ and ‘cholesterol metabolic process’ were enriched in both the outside workers (Fig 3b) and the old queens (Fig 3c). The first common term contained four ‘proteasome subunit’ related genes that were overexpressed in both outside workers and older queens (*Lneg_g05162* BLASTp match proteasome subunit alpha type-5 [*Ooceraea biroi*] / XP_011339939.1, *Lneg_g05171* match proteasome subunit alpha type-6 [*Solenopsis invicta*] / XP_011167410.1, *Lneg_g13954* match proteasome subunit alpha type-1 [*Temnothorax curvispinosus*] / XP_024875312.1, *Lneg_g16342* match proteasome alpha4 subunit [*Drosophila melanogaster*] / NP_525092.1). Similarly, two predicted hormone-sensitive lipases [*Linepithema humile*] associated with the genes *Lneg_g10026* (BLASTp match XP_012229728.1) and *Lneg_g10677 (*XP_012229728.1) were upregulated in both the outside workers and the old queens (Supplementary Table 6).

**Figure 3:**
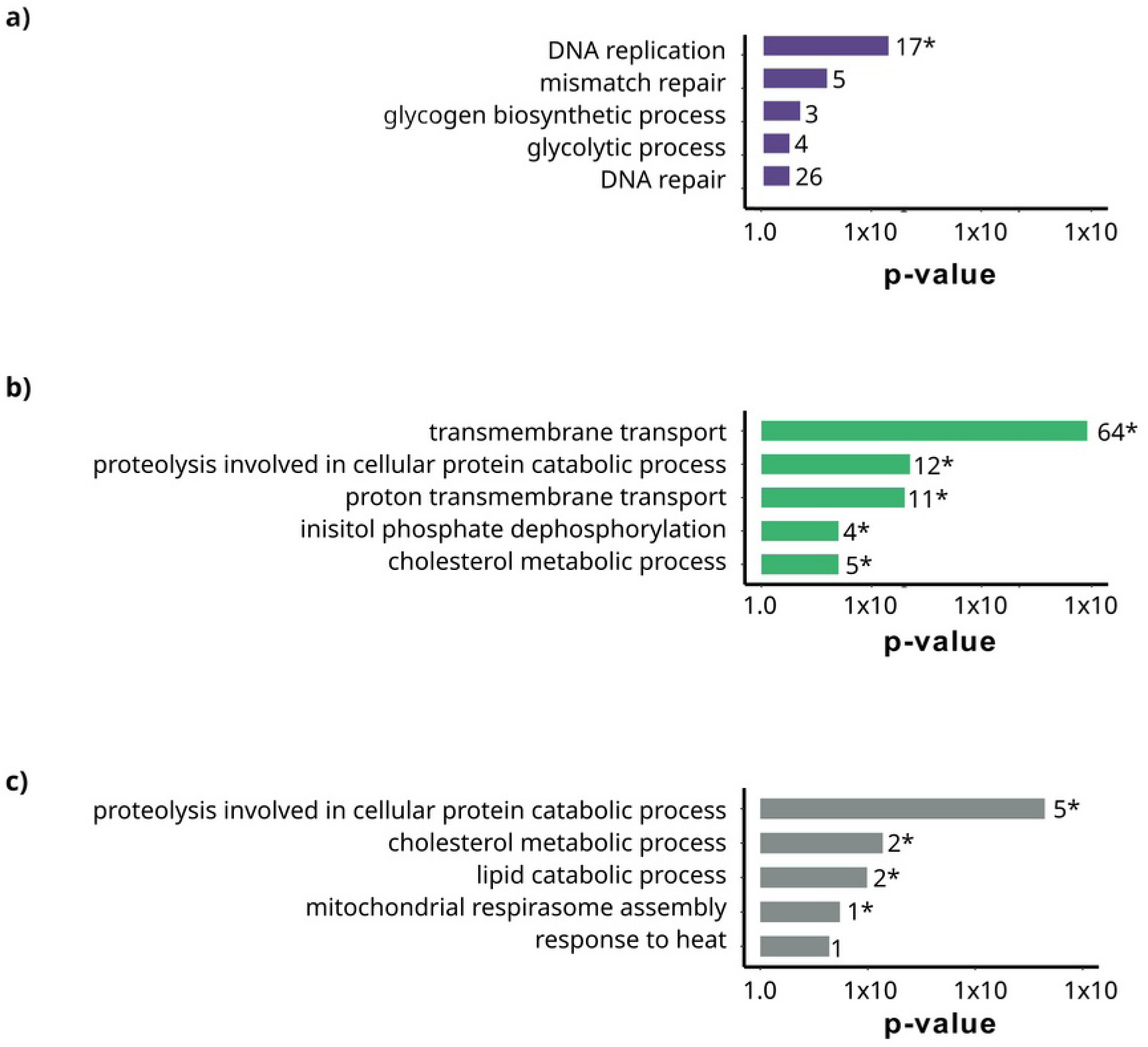
The five most enriched GO terms for Biological Processes based on genes upregulated in younger nursing (inside) *versus* older foraging (outside) workers (a), in foragers vs. nurses (b), and in older queens versus younger queens (c). No genes were upregulated in young queens relative to old queens. GO terms are presented from top to bottom in decreasing order of p-value significance in Fisher’s exact tests after Benjamini-Hochberg correction (* indicates p < 0.05). The number of DEGs associated with each GO Term is given next to each bar (Supplementary Table 6). Bar colours are the same as in Figure 2.

### Ageing and oxidative stress related genes

We found that 66 common orthogroups contained the proteins associated with the 123 candidate genes involved in IIS (Insulin/Insulin Like), TOR (Target of Rapamycin), and Juvenile hormone (JH) signalling pathways in *Drosophila melanogaster*, which have all been implicated in lifespan regulation and ageing in social insects (referred to as ‘TIJ-LiFe’ by Korb et al., 2021). We compared this list with our DEGs (p_adjusted_ < 0.05) for both the workers and the queens and found seven unique orthogroups containing 10 of the 1444 genes upregulated in inside workers. This number did not exceed what would be expected by chance, indicating that TiJ-LiFe genes were not particularly upregulated in inside workers (χ^2^ = 1.4 e^-25^, p = 0.99). This included genes such as four copies of *jhamt*, the TOR complex 1 regulator *gig,* and the gene *thor,* which encodes a translation initiation factor *4-E binding protein,* which is in turn regulated by *tor.* The genes overexpressed in outside workers contained 11 of all 66 orthogroups containing ‘TIJ-LiFe’ genes. They contained 17 out of 1229 DEGs, which is more than expected by chance (χ^2^ = 3.99, p = 0.04). Among these genes were *ImpL2 and* two copies of the gene *NLaz*, all inhibitors of IIS signalling. Only one of the genes that were overexpressed in young queens (*pdk1*) and two genes overexpressed in old queens (one copy of a *Lipase* gene and one copy of a *jhamt*) were ‘TIJ-LiFe’ genes (Supplementary Table 7).

For the analysis of oxidative stress genes, we employed the same method as above and found 31 unique orthogroups containing the 33 genes related to oxidative stress (85 proteins) in *D. melanogaster* and previously tested in other social insects (Kramer et al., 2021). Of these genes, the inside workers had upregulated expression of two copies of *Catalase*, one *Grx5* and one *gfzf* gene. The foraging workers collected outside the nest had upregulated expression of seven oxidative stress-associated genes from seven unique orthogroups. These included: *Grx1*, *MsrA*, *Trx-2 and Trxr2*. None of these oxidative stress genes occurred in the list of queen DEGs (Supplementary table 7).

Finally, we found four matches when we conducted a reciprocal BLASTp of the whole *L. neglectus* proteome against a Vitellogenin database (Kohlmeier et al., 2018). Two genes *Lneg_Vitellogening002* and *Lneg_g15702* were overexpressed in inside workers relative to outside workers (both Dseq2 padj < 0.0001). They encode proteins that showed BLASTp matches with ‘Vitellogenin’ of *Lasius niger* (KMQ94142.1) and ‘Vitellogenin-like protein A’ of *Formica exsecta* (AII96913.1), another ant genus of the same subfamily. Both these vitellogenins have been classified as *Vg-like A* copies by Kohlmeier et al. (2018; Supplementary Table 4). The expression of *Vg-like C* (encoded by the gene *Lneg_g03782*), with a BLASTp match of AII96915.1 with *Vg-like C* in *Formica exsecta*, tended to be higher in outside workers relative to inside workers (Dseq2 padjust = 0.050). The protein of gene *Lneg_g11491,* which matched the ‘vitellogenin-precursor’ of *L. niger,* was classified as the *Conventional-Vg/C-Vg*. DESeq2 did not generate a p-value in workers because of outliers, but there was little overlap in the expression levels of young and old workers, with young workers expressing much more C-Vg than old workers (log_2_fold change = −5.3, SE = 0.8; Supplementary Table 7). *C-Vg* was marginally overexpressed in younger queens relative to older queens, but its expression was not significantly explained by queen age (*Lneg_ g11491 Dseq2* padjust = 0.105) (Supplementary Table 7).

## Discussion

We tested the hypothesis that the molecular physiology of *Lasius neglectus* workers has evolved to become independent of queen presence because these workers are fully sterile and thus represent the true collective soma of their colonies. Our findings support this hypothesis, as the presence of queens did not affect the susceptibility of workers to oxidative stress or alter the transcriptomes of their fat bodies, in contrast to other ant species whose queenless workers often develop their ovaries and become longer lived to raise any remaining brood before the colony collapses (Dijkstra et al. 2007, Negroni et al. 2021). However, for *L. neglectus* we found substantial differences in gene expression and stress resistance between younger nursing workers and older foragers, consistent with workers changing their tasks with age. We found similar differences between younger and older queens, but to a smaller extent.

### Queen loss does not affect worker survival and fat body transcriptomes

Our experiments demonstrated that the removal of queens did not affect the susceptibility of *L. neglectus* workers to paraquat-induced stress, and neither did this treatment alter the transcriptional activity of worker fat bodies. There were, in fact, no genes that differed in expression between workers with and without queens, in sharp contrast with many studies reporting that queen removal triggers a cascade of phenotypic and regulatory changes in the worker caste: In both ants and honeybees, queen loss affects the expression of hundreds of genes, especially genes encoding proteins associated with reproduction and the alleviation of oxidative stress (Alaux et al 2006; Amsalem et al., 2017; Giehr et al., 2020b; Kennedy et al., 2021a; Negroni et al., 2021; Seehuus et al., 2006a; Tsuji et al., 1996). Queenless workers in other ants were also typically longer lived or more stress resistant than workers with queen (Majoe et al., 2021; Choppin, 2022; Negroni et al., 2021), possibly because molecular pathways for egg production also affect body maintenance. For instance, the egg yolk precursor vitellogenin protects honeybee workers from oxidative stress (Münch & Amdam, 2010).

Our finding that queen removal in *L. neglectus* did not affect worker physiology matches the expectation that the high number of queens in unicolonial ants must have relaxed selection for worker responses to queen loss. Similar forms of functional decoupling between colony germline and colony soma individuals may thus have convergently evolved in other ants with a comparable unicolonial and invasive population structure (Helanterä et al., 2009), but this remains to be tested. It is also not entirely clear if queen loss might affect the gene expression in other worker organs, particularly the brain. It would also be of interest to clarify why *L. neglectus* queens still produce queen pheromones (Holman et al. 2013). If these pheromones do not affect worker behaviour, it is tempting to speculate that they have obtained a secondary role in the regulation of queen density and migration across neighbouring nests, so that a rather constant queen-worker ratio can be maintained throughout the entire colony that may extend over many hectares (Espadaler & Rey, 2001).

### Nursing and foraging workers differ in gene expression and susceptibility to stress

We investigated the fat body gene expression patterns of nursing workers collected near the brood (‘inside workers’) and of workers from the foraging arena (‘outside workers’) and found substantial differences, consistent with these workers performing brood care and foraging tasks. One of the clearest patterns was in the expression of genes coding for vitellogenins. These proteins are typically involved in lipid transport into the eggs, but several homologs that arose after a series of duplications of the ancestral ‘conventional’ *Vg* gene have been suggested to also be differentially expressed depending on caste and task (Corona et al 2013; Morandin et al., 2014; Salmela et al., 2016; Wurm et al., 2011). In our *L. neglectus* data, *Vg-like A* was overexpressed in nursing workers while *Vg-like C* was overexpressed in foraging workers. Similar patterns were found in the fat bodies of *Temnothorax longispinosus* ants (Kohlmeier et al., 2018, 2019), a *Diacamma* species (Miyazaki et al., 2021), and in whole-body transcriptomes of *Formica fusca* (Morandin et al., 2019).

Nursing and foraging workers differed in more than just the task they were performing. Although there are no explicit data on the age-specific trajectory of typical worker tasks in *L. neglectus*, it is fair to assume that the workers sampled outside the nest were older than those engaged in tasks inside the nest. This is typically the case across ant species and in social insects in general (Bernadou et al., 2015; Giraldo & Traniello, 2014; Pamminger et al., 2014; Ravary et al., 2007). The main pattern of what is sometimes referred to as temporal or age polyethism is that workers take up brood care shortly after hatching and become engaged in hazardous tasks such as foraging and defence as they get older. The results of our gene expression analysis confirmed this universal expectation as the two categories of workers varied in the expression of over 2000 genes, among them a number of genes typically involved in ageing. Relative to younger nursing workers, the workers operating outside the nest overexpressed proteolysis genes such as proteasome subunits. This is consistent with older workers accumulating carbonylated proteins, which are then recycled by proteasomes (de Verges & Nehring 2016, Kramer et al 2021).

When we investigated the resistance to oxidative stress, we found that younger nursing workers survived better than older foraging workers. This difference was visible both in the workers exposed to paraquat-induced oxidative stress, and in the control, workers treated with water. However, more than half of the control workers died in the 14-day survival experiment, suggesting that experimental isolation in small sub-colonies followed by experimental manipulation every other day was very stressful by itself. We could not disentangle the potential effects of task and age and can therefore not exclude that outside workers died earlier because mortality is naturally higher in older workers. However, *L. neglectus* workers living longer than seven months have been recorded (Espadaler & Rey, 2001), making it unlikely that natural age-specific mortality alone can explain the high death rate of control workers. Our results thus continue to suggest that the older *L. neglectus* workers collected outside the nest were more susceptible to stress than the younger nursing workers, consistent with earlier findings in leaf-cutting ants (Majoe et al., 2021) and honeybees (Alqarni et al., 2019; Amdam et al., 2005; Seehuus et al., 2006).

### Age-related effects on queen gene expression

Fat bodies of older *L. neglectus* queens differed from those of young queens in the expression of 165 genes, which is a considerably smaller difference than reported for queens of other ant species (Negroni et al., 2019; von Wyschetzki et al., 2015). In our study all factors other than queen age were kept constant, in contrast to previous studies where young queens had not yet completed independent colony establishment and thus lived in a different social environment than older queens (e.g. Negroni et al., 2019). This suggests that previous differences in age-related gene expression of ant queens may have been overestimates. Comparing our queen and worker results also had its challenges because we did not know the individuals’ exact ages relative to average caste-specific lifespans. Our sampling strategy also differed between the castes, as queen samples consisted of single individuals while multiple workers were pooled per replicate so that individual variation was averaged out. Nevertheless, we could detect interesting consistencies in gene expression of old queens and old workers, such as GO terms relating to proteolysis and cholesterol metabolism. This suggests that both older queens and older workers need increased efforts to keep their enzymes and cell membranes functional.

Although we did not know the exact age of the older queens, there was no indication that young queens were healthier than old queens or that the old queens in our experiments were close to the natural end of their lives. In the ant *Cardiocondyla obscurior*, age does not strongly affect gene expression until the queens are about to die (Jaimes-Nino et al., 2022; von Wyschetzki et al., 2015). The typical excess of queens in invasive unicolonial *L. neglectus* populations may have implied selection for faster aging and shorter life span, as is generally the case in polygynous ants (Keller & Genoud, 1997). However, the old and young queens in our study did not appear to differ in fecundity as their ovaries were of similar size (Supplementary Figure 1) and fat body transcriptomes did not differ in the expression of fecundity genes such as C*onventional vitellogenin.* Our results thus support the general view that social insect queens do not age linearly over their reproductive lifetimes, but that they deteriorate very rapidly toward the end of their lives, both reproductively and somatically (Elsner et al., 2018; Jaimes-Nino et al., 2022).

### Conclusion

As in many social wasps and bees, ant workers can develop their ovaries and produce male offspring from unfertilized eggs (Wenseleers & Ratnieks 2006). The likelihood of such worker reproduction is much higher after the death of a single colony queen when worker-queen conflict over male parentage has disappeared. This implies that fertility pheromones that signal queen presence have remained deeply conserved, not only to regulate colonial homeostasis but also to reset worker molecular physiology when worker reproduction becomes a common interest for all colony members (Ratnieks et al 2006). Worker ovaries have become rudimentary in some ant species but selection on queens may well have retained the production of queen pheromones for other reasons than communication queen presence to the worker caste.

The main result of our study is clear and important: The very large number of queens in natural colonies of a unicolonial ant species may cause the somatic worker caste to evolve complete physiological independence from queens because their presence can be taken for granted. We expect that our results apply more generally across invasive unicolonial ants. It seems well-established that the single evolutionary origin of sociality in the ancestral ant required strict lifetime monogamy of a single colony-founding queen (see e.g. Boomsma, 2022). However, the unique higher-level germline function of queens – analogous to higher metazoan germline functionality – has often been secondarily elaborated, particularly by the repeated convergent evolution of the adoption of daughter queens into existing colonies (secondary polygyny, Boomsma et al. 2014). In a subset of these ant lineages, extreme degrees of polygyny produced the unicolonial population structures that preadapted some ant species to becoming invasive pests (Helanterä et al. 2009). The ancestral model of unitary colony growth thus gradually gave way to its opposite, a form of modular growth analogous to how plants and fungal mycelia develop. In modular organisms there is no sequestration of an initially dormant germline shortly after a zygote develops, but exclusive somatic growth until multiple germline organs are formed as flowers or spore-bearing bodies. Further work to investigate whether this analogy applies generally to most if not all highly polygynous and unicolonial ants thus seems of interest for enhancing our general understanding of social evolution.

## Ethics

We did not require any licenses to collect and transport these ants from the Jena Botanical Garden in Germany to our laboratory in Freiburg. We followed the guidelines of the Study of Animal Behaviour and the legal and institutional rules.

## Data accessibility

The raw data behind the Figures are provided in the electronic supplementary material. The genome assembly and annotations used for *L. neglectus* have been generated as part of the Global Ant Genomics Alliance (GAGA) project; these can be accessed upon request to the corresponding author and the GAGA consortium.

## Author contributions

VN, SF, RL and MM designed the experiment. Ants have been collected by VN and MM. JV, ZX and LS from the GAGA consortium, co-coordinated by JJB, conducted the genome assembly and JV and ZX annotated the genes. NS, supervised by SF and co-supervised by MM, cleaned and primed the transcriptomic data for mapping and the analyses that MM conducted. VN, RL and SF helped MM in the statistical analyses and interpretation. MM wrote a first draft and all authors commented on it. MM, VN, SF, RL, and JJB finalized the manuscript based on these comments. VN, SF, and RL jointly supervised the experimental parts of the project.

## Acknowledgements

We thank the Jena Botanical Garden for allowing us to collect ants on their property. We are indebted to Julius Rombach and Anna Waffender for the upkeep and maintenance of the colonies and for conducting the oxidative stress experiments. Thanks to Anna Lenhart for her insights during the writing phase and Barbara Feldmeyer for help with the *vitellogenin* classification.

## Supplementary Material

**Supplementary figure 1:**
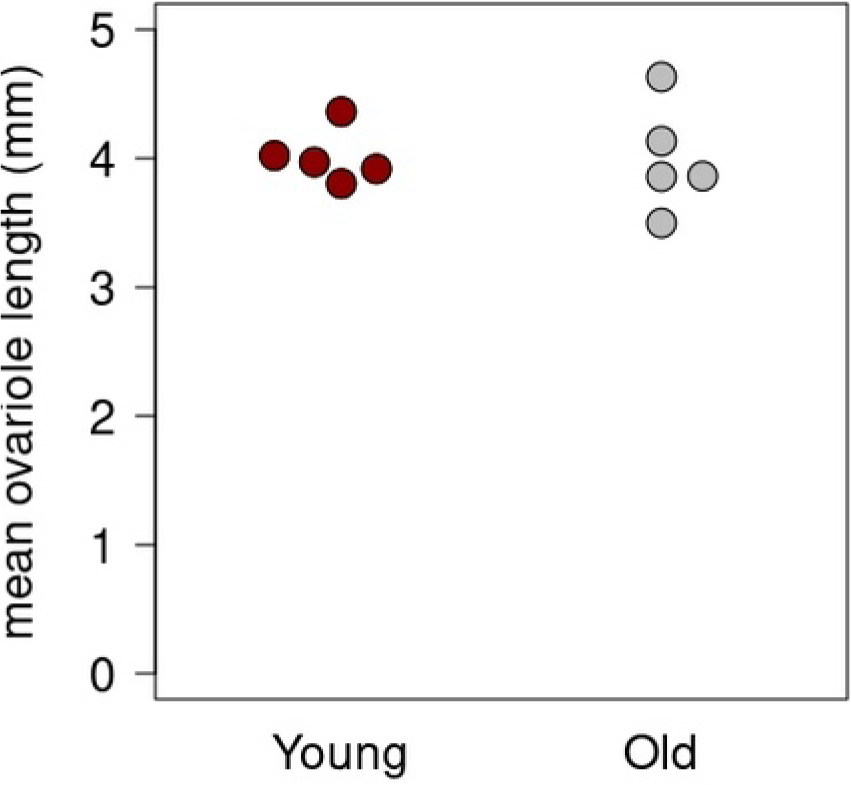
Box plots mean ovariole length measured in five young and five old *Lasius neglectus* queens. Ovariole length of older queens was more variable but the mean length did not differ from the mean of young queens (U-test n = 10, U = 11, p = 0.84).

**Supplementary Table 1.** Information on all worker samples used for RNA-seq analysis.

| **Sample** | **Colony** | **Caste** | **Queen** | **Age** |
| --- | --- | --- | --- | --- |
| Lne22NQIn1 | Lne22 | Worker | Queenless | Young |
| Lne22NQIn2 | Lne22 | Worker | Queenless | Young |
| Lne23NQIn1 | Lne23 | Worker | Queenless | Young |
| Lne23NQIn3 | Lne23 | Worker | Queenless | Young |
| Lne24NQIn1 | Lne24 | Worker | Queenless | Young |
| Lne26NQIn1 | Lne26 | Worker | Queenless | Young |
| Lne26NQIn2 | Lne26 | Worker | Queenless | Young |
| Lne22NQOut | Lne22 | Worker | Queenless | Old |
| Lne23NQOut1 | Lne23 | Worker | Queenless | Old |
| Lne23QNQOut2 | Lne23 | Worker | Queenless | Old |
| Lne24NQOut1 | Lne24 | Worker | Queenless | Old |
| Lne26NQOut1 | Lne26 | Worker | Queenless | Old |
| Lne26NQOut2 | Lne26 | Worker | Queenless | Old |
| Lne22QIn2 | Lne22 | Worker | Queenright | Young |
| Lne22QIn3 | Lne22 | Worker | Queenright | Young |
| Lne23QIn1 | Lne23 | Worker | Queenright | Young |
| Lne23QIn2 | Lne23 | Worker | Queenright | Young |
| Lne23QIn3 | Lne23 | Worker | Queenright | Young |
| Lne24QIn1 | Lne24 | Worker | Queenright | Young |
| Lne24QIn2 | Lne24 | Worker | Queenright | Young |
| Lne26QIn1 | Lne26 | Worker | Queenright | Young |
| Lne22QOut1 | Lne22 | Worker | Queenright | Old |
| Lne22QOut2 | Lne22 | Worker | Queenright | Old |
| Lne23QOut1 | Lne23 | Worker | Queenright | Old |
| Lne23QOut2 | Lne23 | Worker | Queenright | Old |
| Lne24QOut1 | Lne24 | Worker | Queenright | Old |
| Lne26QOut1 | Lne26 | Worker | Queenright | Old |
| Lne26QOut2 | Lne26 | Worker | Queenright | Old |
| Lne26QOut3 | Lne26 | Worker | Queenright | Old |

**Supplementary Table 2.**
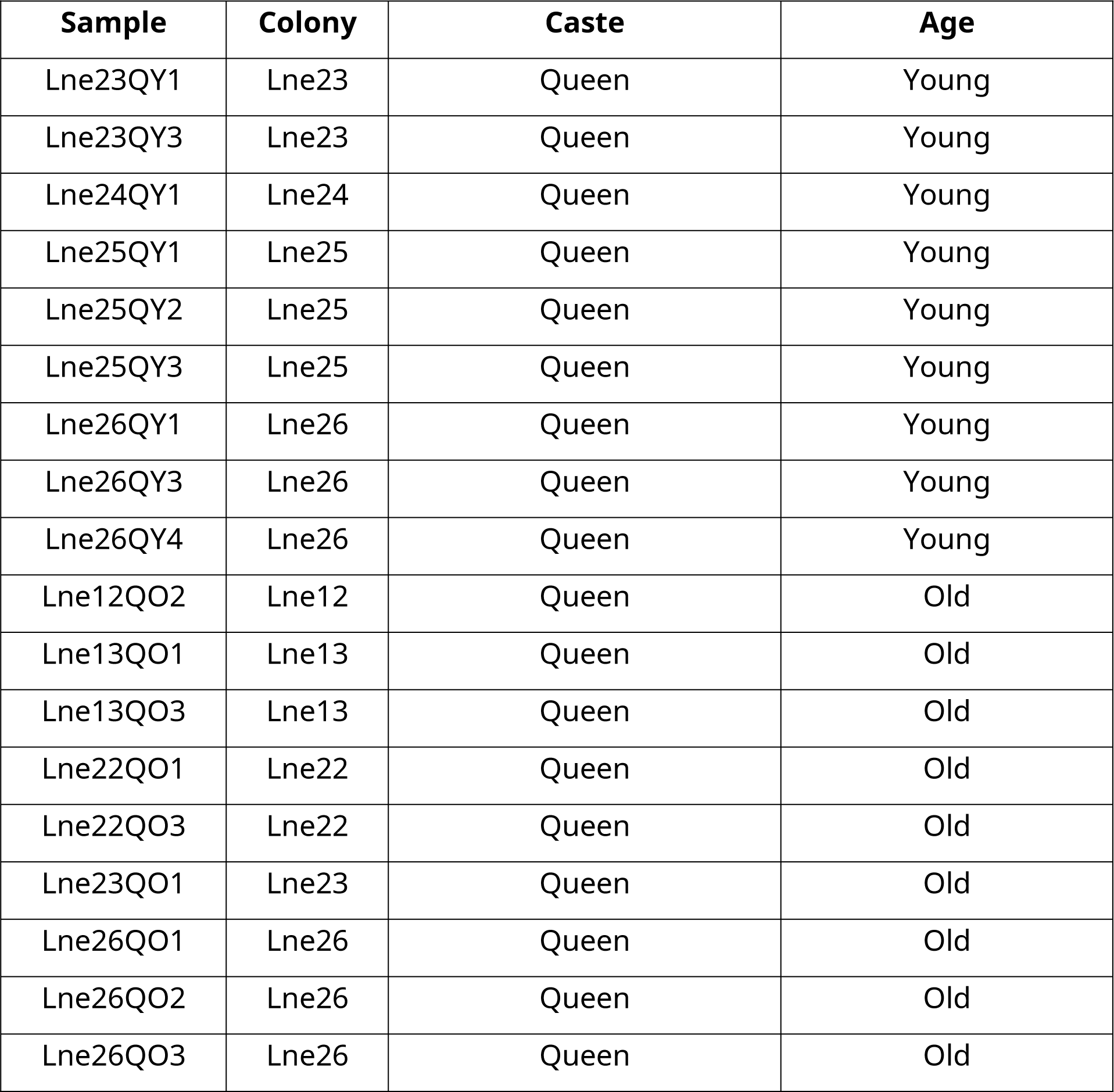
Information on all queen samples used for RNA-seq analysis.

Supplementary Table 3: Genome statistics Supplementary Table 4: BlastP hits

Supplementary Table 5: Data and statistics for the oxidative stress experiment

Supplementary Table 6: Differentially expressed genes and statistics

Supplementary Table 7: Candidate genes (TiJ-LiFe, oxidative stress, vitellogenins)

